# Oligosaccharide production and signaling correlate with delayed flowering in an *Arabidopsis* genotype grown and selected in high [CO_2_]

**DOI:** 10.1101/2023.06.19.545630

**Authors:** Hannah Kinmonth-Schultz, Stephen Michael Walker, Kerem Bingol, David W. Hoyt, Young-Mo Kim, Lye Meng Markillie, Hugh D. Mitchell, Carrie D. Nicora, Ronald Taylor, Joy K. Ward

## Abstract

Since industrialization began, atmospheric CO_2_ ([CO_2_]) has increased from 270 to 415 ppm and is projected to reach 800-1000 ppm this century. Some *Arabidopsis* ecotypes delayed flowering in elevated [CO_2_] relative to current [CO_2_], while others showed no change or accelerations. To predict genotype-specific flowering behaviors, we must understand the mechanisms driving flowering response to rising [CO_2_]. [CO_2_] changes alter photosynthesis and carbohydrates in C_3_ plants. Plants sense carbohydrate levels and exogenous carbohydrate application influences flowering time and flowering transcript levels. We asked how organismal changes in carbohydrates and transcription correlate with changes in flowering time under elevated [CO_2_]. We used a genotype (SG) of *Arabidopsis* that was selected for high fitness at elevated [CO_2_] (700 ppm). SG delays flowering under elevated [CO_2_] (700 ppm) relative to current [CO_2_] (400 ppm). We compared SG to a closely related control genotype (CG) that shows no [CO_2_]- induced flowering change. We compared metabolomic and transcriptomic profiles in these genotypes at current and elevated [CO_2_] to assess correlations with flowering in these conditions. While both genotypes altered carbohydrates in response to elevated [CO_2_], SG had higher levels of sucrose than CG and showed a stronger increase in glucose and fructose in elevated [CO_2_]. Both genotypes demonstrated transcriptional changes, with CG increasing genes related to fructose 1,6-bisphosphate breakdown, amino acid synthesis, and secondary metabolites; and SG decreasing genes related to starch and sugar metabolism, but increasing genes involved in oligosaccharide production and sugar modifications. Genes associated with flowering regulation within the photoperiod, vernalization, and meristem identity pathways were altered in these genotypes. Elevated [CO_2_] may act through carbohydrate changes to influence transcription in both genotypes and delayed flowering in SG. Changes in the oligosaccharide pool may contribute to delayed flowering in SG. This work extends the literature exploring genotypic-specific flowering responses to elevated [CO_2_].

## Introduction

Our planet is experiencing an increase in the concentration of atmospheric carbon dioxide ([CO_2_]) that is unprecedented on the scale of evolutionary time. Atmospheric [CO_2_] has increased from 270 ppm at the onset of the industrial age to a current value of 410 ppm due to fossil fuel combustion, and it is projected to increase to 700 ppm in the next 100 years [1]. This has implications for agricultural and ecological systems as plants have experienced relatively low [CO_2_] over the last several million years, with minimums of 180 ppm occurring during peak glacial periods as recently as 20,000 years ago. One critical effect is changes in the timing of plant phenological events, including timing of peak flowering, which depending on the direction of change, may alter pollinator interactions [2–6], but see [7], or increase exposure to stressful climactic events such as spring frosts or summer droughts [2,8– 11].

Much focus has been on warming temperatures occurring concomitantly with [CO_2_] change (1.09 °C rise since 1850 [1]); however, recent evidence suggests that [CO_2_] change contributes both independently and interactively with temperature to influence flowering time. For example, work with several field-collected *Arabidopsis thaliana* accessions isolated the separate and interactive effects of temperature and [CO_2_] rise since the onset of the industrial era [12]. This work demonstrated that temperature and [CO_2_] changes interacted to accelerate flowering in modern conditions compared to pre-industrial conditions. [CO_2_] also independently influences flowering as a comprehensive review demonstrated that 57% of the wild species and 62% of the crop species tested exhibited changes in flowering times when grown at elevated [CO_2_] (projected for year 2100) versus 350-380 ppm [CO_2_] (modern levels) [13].

Currently, the patterns of [CO_2_]-induced flowering time shifts are far from predictable as the magnitude and direction of changes in response to [CO_2_] vary both intra- and interspecifically, and are modulated not only by temperature, but other environmental variables. Tested species showed accelerations, no change, or delays in flowering times in response to [CO_2_] changes alone [13], and parallel ranges of response are observed within species. For example, in our work, two *Arabidopsis* strains from the same parental cross differed strikingly in their [CO_2_]-induced flowering time responses. Although grown in the long-day conditions needed to induce flowering, a strain selected for high fitness, as measured by seed set, over successive generations in high [CO_2_] (selected genotype, SG) delayed flowering by 10 d or more at elevated [CO_2_] (700ppm) relative to 380 ppm. The control genotype (CG), that arose from random selection of individuals over successive generations, did not alter its flowering phenotype [14,15]. Similarly, near-isogenic soybean lines from the same genetic background differed in [CO_2_] sensitivity, and the direction of change was modulated both by genotype and daylength [16]. Lines with dominance in single photoperiod genes accelerated flowering at elevated [CO_2_] (560 ppm) in longer daylengths but delayed in shorter daylengths compared to modern [CO_2_], while lines recessive across all photoperiod genes delayed flowering at elevated [CO_2_] under all daylengths.

The mechanisms behind [CO_2_]-induced flowering time changes and the variation in this response are unknown; however, alterations in the carbohydrate compositions in plant tissue likely contribute to this shift. Carbon dioxide concentration changes lead to marked changes in the rate of carbon accumulation through photosynthesis and in insoluble and soluble carbohydrates, downstream metabolites, and the carbon-to-nitrogen ratio across species [17]. Further, alterations in both photosynthesis and carbohydrates composition have been linked to flowering. For example, *Lolium tremulentum* delayed flowering when treated with DCMU (3-(3,4-dichlorophenyl)-1,1- dimethylurea), a photosynthesis inhibitor [18], and floral induction coincides with a flux of carbohydrates from the leaves to the shoot apex [19–23]. Additionally, sucrose application in the growth media delayed flowering in *Arabidopsis*, a facultative long-day plant, with delay positively correlated to sucrose concentration [24]. The delay was caused by a longer vegetative phase, resulting in more leaves before flowering and was coupled with lower expression of the floral inducer genes *FLOWERING LOCUS T* (*FT*) and *LEAFY* (*LFY*). This behavior was similar to the delays in the soybean isolines at elevated [CO_2_], which were associated with a higher number of nodes on the main stem, although carbohydrate content was not tested in that work [16].

Work in a wide range of organisms has highlighted a connection between carbohydrate or nutritional variation, downstream metabolic shifts, and global or targeted gene transcriptional regulation, providing possible specific mechanisms through which [CO_2_] may influence developmental change. For example, in *Arabidopsis,* trehelose-6-phosphate (T6P) is responsive to sucrose levels and influences the expression levels of *FT* [25]. The authors proposed that T6P plays a role in signaling when carbohydrate reserves are sufficient to support the energy demands of reproduction. Additionally, a reversable post-translational sugar modification, O-linked B-N- acetylglucosaminylation (O-GlcNAcylation), is involved in the regulation of several transcriptional and epigenetic regulators including RNA polymerase II (Pol II) and Polycomb group (PcG) proteins across a variety of organisms [26–28]. It is linked to nutrition and carbohydrate-related diseases in humans such as *in utero* epigenetic responses in mothers exposed to famine or having Type-2 diabetes [26,29]. In plants, O-GlcNAcylation modulates the function of DELLA family proteins as well as expression of key flowering repressor gene *FLOWERING REPRESSOR C* (*FLC*) [30,31]. Both DELLAs and PcG proteins are involved in numerous endogenous and exogenous signaling pathways controlling plant environmental perception, development, and flowering time (e.g. [32–34]). Finally, the serine and glycine pools, downstream products of the Calvin cycle and glycolysis, shunt carbon through one-carbon metabolism to affect epigenetic regulation and such processes as cancer oncogenesis, further linking metabolism and nutrition to cell regulation [35,36]. In plants, the same process also acts downstream of the photorespiratory pathway to affect a range of processes including methylation and auxin synthesis and appears important for plant development and environmental response [37]. All three processes offer intriguing mechanisms that could not only influence [CO_2_]-induced flowering time changes but explain how [CO_2_]-driven responses are modulated by interactions with other environmental variables. Additionally, both standing levels of carbohydrates [38,39] and photosynthesis [40,41] and the degree to which they respond to different [CO_2_] differ between and within species [41–44] even after just eight days of growth in elevated [CO_2_] [45], thus suggesting a possible mechanism of intraspecific variation in response to [CO_2_].

Compared to environmental processes such as daylength and temperature-mediated flowering time control [34,46–48], our knowledge of the molecular processes governing [CO_2_]-driven flowering time shifts and the variation in that response is very much in its infancy (e.g. [15]). To begin to understand the mechanisms of [CO_2_]- induced flowering time changes and the variation in this response, we utilize the CG SG system of *Arabidopsis* described above [15]. The CG SG system we describe above is a useful model, because as observed with *Arabidopsis* exposed to increasing concentrations of sucrose and soybean isolines exposed to elevated [CO_2_] [16,24], the delay in flowering in SG appears not only to be influenced by changes in the *rate* of growth and development, as would be expected from increased carbon acquisition through photosynthesis, but by alteration of the *developmental stage* (plant size and/or leaf number) at which plants flower. In our work, although SG developed more rapidly at elevated [CO_2_] compared to current [CO_2_], it flowered later because it produced more leaves before transitioning to the reproductive stage [15]. Conversely, CG flowered at a similar leaf number in both [CO_2_]. This delay was correlated with prolonged high expression of the cold-responsive, flowering repressor gene *FLOWERING LOCUS C* (*FLC*) that acts upstream of *FT* only in SG grown in elevated [CO_2_] [15]. In fall-germinating, winter-annual variants of *Arabidopsis*, *FLC* stays high until it is repressed by prolonged cold (=vernalization) [49] likely so that *FT* rises and flowering occurs appropriately in warm, spring conditions. In subsequent work, we confirmed that downregulation of *FLC* through vernalization restored early flowering in SG in elevated [CO_2_] [50]. However, the upstream mechanisms governing the response of *FLC* to [CO_2_] are unknown. Additionally, another flowering pathway may also be altered by [CO_2_] change, as *LFY* was altered in both SG and CG, at least partially independently of *FLC* [15,50].

Here, by assessing the correlations between transcriptional and carbohydrate profiles and their relationship to known flowering time regulators, we aimed to evaluate through what metabolic pathways [CO_2_] changes may be acting to influence flowering genes and flowering and how these processes differ between the genotypes at different [CO_2_]. Thus, we harvested SG in current [CO_2_], just prior to the visible transition to reproduction and SG in elevated [CO_2_] at the analogous leaf number, but well before the reproductive transition. For comparison, we harvested CG in current and elevated [CO_2_] just prior the reproductive transition as well. We specifically ask, how are carbohydrate profiles altered by [CO_2_] across genotypes before flowering or at the analogous developmental stage in SG at elevated [CO_2_], what transcriptional pathways correlate with the carbohydrate changes, and how does the [CO_2_] response in SG vary from that of CG.

## Results

### Increased glucose and fructose correlates with elevated-[CO_2_]-induced flowering delay in Selected Genotype

To assess carbon acquisition capacity of both the CG and SG lines at different [CO_2_] and to determine the mechanisms responsible for genotype-specific flowering behaviors and the delay in flowering in SG, we compared primary carbohydrates—glucose, fructose, and sucrose—in CG just prior to production of a visible flowering stem (bolt) in current (380 ppm) and elevated (700 ppm) [CO_2_], SG just prior to flowering at current [CO_2_], and SG grown in elevated [CO_2_] at the analogous developmental stage (leaf number), which was several days before it would transition to flowering (**Fig. 1**). To do so, we combined data from two experimental replicates that were detected through different methods (GCMS and NMR). To better compare across the two datasets and to be consistent with treatment of other data in this study, we transformed the data using centered-log ratio, then used ANOVA to assess effect of genotype, [CO_2_] level, their interaction, and included replicate as a covariate. All three carbohydrates showed a clear effect of replicate; however, after variation for replicate was accounted for, glucose showed a significant [CO_2_] effect (*p* < 0.001), fructose showed both a significant genotype and [CO_2_] effect (*p* < 0.01, 0.001, respectively), and sucrose showed an effect of genotype (*p* < 0.0001). The genotype by [CO_2_] interaction was not significant across the three carbohydrates. To assess the treatment groups driving these differences, *post hoc* analysis revealed that SG had overall higher sucrose levels, with SG plants grown in 380 and 700 ppm being significantly higher than CG grown in 700 ppm (*p* < 0.01, 0.005, respectively), SG also had higher glucose levels driven by SG in 700 ppm being significantly higher than CG grown in 380 ppm (*p* < 0.05) (**Table 1**). Fructose showed the reverse, with SG in 380 ppm being lower than both CG in 380 and 700 ppm (*p* < 0.05, 0.0001, respectively). However, the effect of [CO_2_] on glucose and fructose appears driven by SG. In both cases, SG grown in 700 ppm had higher levels than SG grown in 380 ppm (*p* < 0.01, 0.05, respectively), while the difference between [CO_2_] treatments in CG was not significant (**Table 1**). In sum, SG appears to accumulate more glucose and sucrose relative to CG, especially in elevated [CO_2_] when flowering in SG is delayed, and SG accumulates more glucose and fructose at 700 ppm [CO_2_] than at 380 ppm, again correlating to a delay in flowering. The increase in primary carbohydrates in SG is consistent with previous observations in *Arabidopsis* as reviewed in [17]; as are the differences among genotypes in their standing levels and responses to [CO_2_] [38,39,45]. Thus, carbon accumulation in the form of glucose and fructose may contribute to the delay in flowering in SG at elevated [CO_2_]. More work is needed to determine whether higher sucrose concentrations are indicative of broader genotype sensitivity to [CO_2_] in terms of phenological shifts.

**Figure 1.**
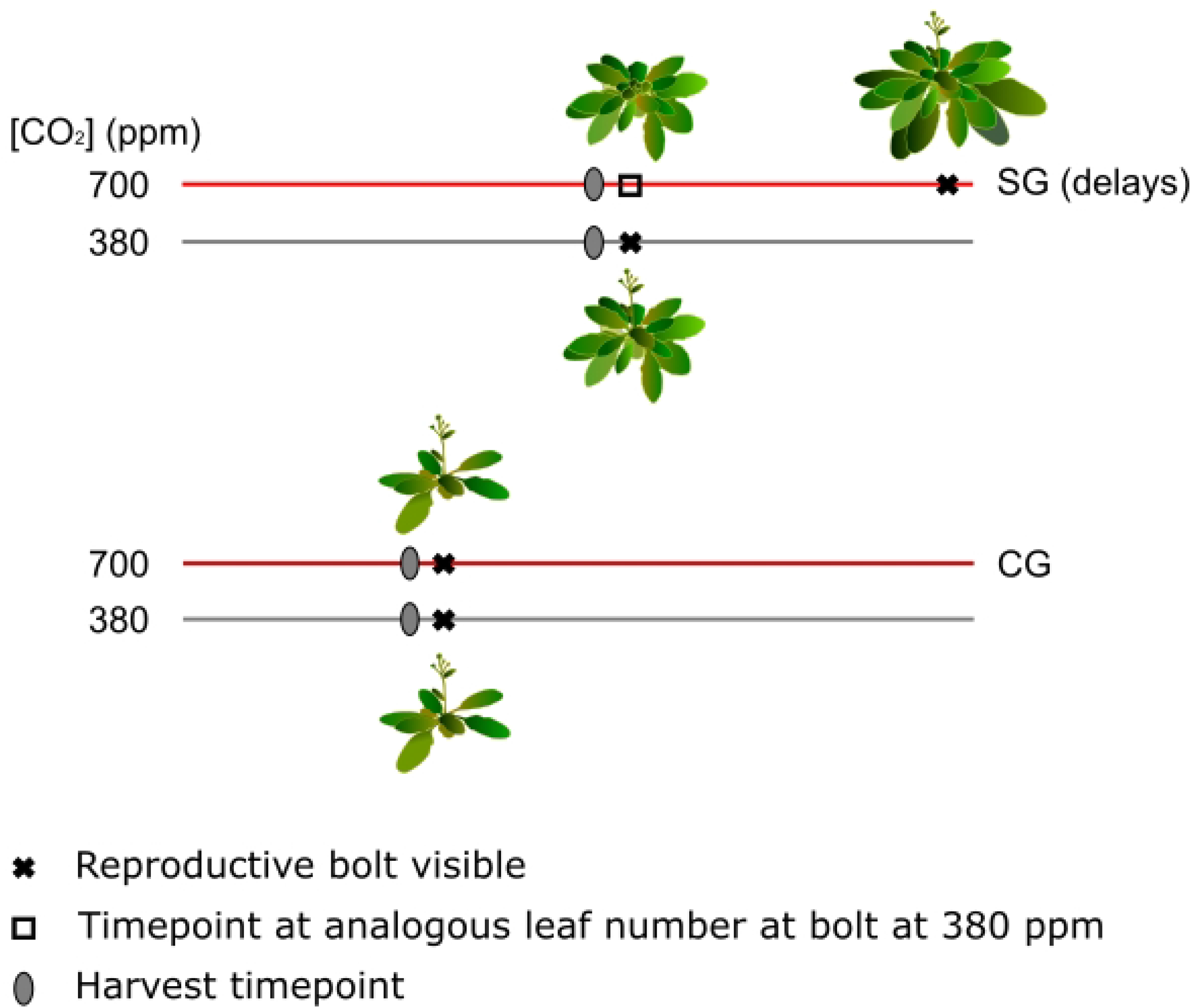
Experimental set up. The Control and Selected Genotypes (CG and SG) were grown both at ‘current’ atmospheric [CO_2_] (380 ppm) and projected future [CO_2_] (700 ppm). Plants were harvested just prior to visible emergence of the reproductive stem (=bolt) for CG at 380 and 700 ppm, and for SG at 380 ppm. For SG at 700 ppm, plants were harvested at the analogous leaf number to SG grown at 380 ppm. Red and grey bars represent time. Leaf numbers shown are approximate. Bolt heights are exaggerated for visibility.

**Table 1:** Results of Tukey’s Honest Significant Difference comparison for primary carbohydrates fructose, glucose, and sucrose. *P* value adjusted for multiple comparisons and upper and lower 95% confidence interval bounds are shown. **diff** = difference between the two treatments being compared in each row.

|  | Comparison | diff | lwr | upr | *p* adj |
| --- | --- | --- | --- | --- | --- |
| **Glucose** | SG:380-CG:380 | -0.0424 | -0.3292 | 0.2445 | 0.9798 |
|  | CG:700-CG:380 | 0.1804 | -0.0986 | 0.4594 | 0.3297 |
|  | SG:700-CG:380 | 0.3272 | 0.0403 | 0.6140 | **0.0191** |
|  | CG:700-SG:380 | 0.2228 | -0.0604 | 0.5059 | 0.1725 |
|  | SG:700-SG:380 | 0.3695 | 0.0786 | 0.6604 | **0.0072** |
|  | SG:700-CG:700 | 0.1468 | -0.1364 | 0.4299 | 0.5249 |
| **Fructose** | SG:380-CG:380 | -0.4555 | -0.8331 | -0.0779 | **0.0118** |
|  | CG:700-CG:380 | 0.2286 | -0.1386 | 0.5959 | 0.3632 |
|  | SG:700-CG:380 | 0.0050 | -0.3727 | 0.3826 | 1.0000 |
|  | CG:700-SG:380 | 0.6842 | 0.3114 | 1.0569 | **0.0000** |
|  | SG:700-SG:380 | 0.4605 | 0.0775 | 0.8434 | **0.0121** |
|  | SG:700-CG:700 | -0.2237 | -0.5964 | 0.1491 | 0.3960 |
| **Sucrose** | SG:380-CG:380 | 0.5621 | -0.1407 | 1.2650 | 0.1612 |
|  | CG:700-CG:380 | -0.3468 | -1.0304 | 0.3368 | 0.5429 |
|  | SG:700-CG:380 | 0.6542 | -0.0486 | 1.3571 | 0.0771 |
|  | CG:700-SG:380 | -0.9089 | -1.6028 | -0.2151 | **0.0053** |
|  | SG:700-SG:380 | 0.0921 | -0.6208 | 0.8049 | 0.9863 |
|  | SG:700-CG:700 | 1.0010 | 0.3072 | 1.6949 | **0.0018** |

**Bold, italicized text in p adj.** column shows significant comparisons.

### Selected and control genotypes vary in their transcriptional responses to [CO_2_]

To further determine the mechanisms responsible for different flowering behaviors between CG and SG, we assessed whether transcriptional patterns, as detected through RNAseq and aligned to the Araport 11 reference genome, were altered between the genotypes and within the genotypes between [CO_2_], using the same comparisons as with the primary carbohydrates, above. Originally, we detected 31,556 unique transcript identifiers across CG and SG. After removing poor-quality or low-count transcript identifiers, 16,472 transcript identifiers remained. We transformed the data using the centered-log ratio (clr) including all transcript counts within each sample in the denominator, as is recommended for compositional data [51,52], then calculated the Aitchison distances across all samples [53]. Note that as each transcript is centered relative to the geometric mean of all transcripts in that sample, values reported are relative to this within-sample control. At this stage, we compared samples based on their distances and found that one sample each from CG and SG grouped apart from the other samples in their strains (**Fig. S1**). These samples were removed as outliers and the distances recalculated. We, then, compared the relationships among the remaining sample distances using Principal Components Analysis (PCA) and permuted multivariate ANOVA (PERMANOVA), with the latter assessing the effects of genotype, [CO_2_], and their interaction. Principal Component 1 (PC1) explained 65.6% of the variation, and this was largely driven by genotype (**Fig. S2a**). No clear pattern based on genotype or [CO_2_] treatment emerged across PC2, which explained 11.1% of the variation; however, SG showed a clear separation between [CO_2_] treatments relative to CG across PC3, which explained 5.3% of the variation (**Fig S2b**). Consistent with this pattern, PERMANOVA revealed a significant genotype effect and genotype-by-[CO_2_] interaction (*p* < 0.001, 0.05, respectively).

To assess the transcripts driving these patterns, we conducted differential expression analysis of each identified transcript. For these, we assessed whether there was an effect of genotype and [CO_2_], and report those with an effect size greater than ±1 [52]. Between genotypes, 3616 fit this condition (**Fig. 2a-b, Table S1**). For, [CO_2_] only 48 genes showed an effect size greater than ±1, with 39 of those showing a decrease relative to the internal control in elevated [CO_2_] (**Table S1**). We used functional annotation clustering to determine the probable function of these altered transcripts (**Table 2 & S2a-d**). This process pulls annotation terms from multiple resources and clusters those terms based on whether they overlap in the genes used to call those terms. For those genes showing strong effect sizes across genotypes and being higher in SG than CG relative to the internal control, the top five functional clusters were: serine/threonine and protein kinase activity; transmembrane or membrane components; calcium-binding region, serine/threonine kinase, peptidyl-serine phosphorylation; signal transduction; ADP and DNA binding, leucine-rich repeats; and ankyrin repeat and PGG domains. For those genes lower in SG than CG, the top five functional clusters were: chloroplast related; ribosome related; chloroplast thylakoid lumen related; ribosomal RNA-binding related; and lipid biosynthesis and metabolism related. For the effect of [CO_2_], when analyzed across genotype, no significant functional annotations emerged (**Table S2d**).To better understand the transcripts driving the significant interaction observed in the PERMANOVA as genotype was such a strong determinant of differences in transcript profiles, we next assessed the effect of [CO_2_] for each genotype individually (**Fig. 2c-f, Table, S1 & S2a,e-h**). CG showed 995 transcripts with a large effect size with most increasing from 380 to 700 ppm [CO_2_], while SG showed 1273 transcripts with a large effect size with most decreasing from 380 to 700 ppm [CO_2_]. For those increasing in CG (**Fig. 2c**, **Table 2, S1 & S2a,e**)., the top five functional annotation clusters were: transit peptide, chloroplast thylakoid membrane; PSI, PSII, chlorophyll a/b, chloroplast, magnesium binding; membrane, transmembrane; glycolysis/gluconeogenesis, biosynthesis of amino acids/antibiotics; cytochrome b5 heme-binding. Each of these clusters also included the annotation terms ‘plastid’, ‘chloroplast thylakoid’, ‘chloroplast envelope’, ‘Chlorophyll-a 5’, and ‘Chlorophyll-a 5’, respectively, potentially indicating that much of the increased transcript activity within CG at elevated [CO_2_] was associated with photosynthetic structures. For those decreasing in CG (**Fig. 2d**, **Table 2, S1 & S2a,f**)., the top five clusters were: ATP and nucleotide binding; microtubule motor protein activity; nucleus, sequence-specific DNA binding, and transcription regulation; cell division and mitosis; and zinc and metal binding. For those transcripts increasing in SG (**Fig. 2e**, **Table 2, S1 & S2a,g**)., the top five clusters were: Golgi related and glycosyl and hexosyl transferase activity; small GTP binding and GTPase-mediated signal transduction; cytoskeleton and microtubule associated; cell wall organization; and IQ motif and calmodulin binding. For those decreasing in SG (**Fig. 2f**, **Table 2, S1 & S42a,h**)., the top five clusters were: Chloroplast and transit peptide; carbon metabolism and fixation, biosynthesis and metabolic pathways; thylakoid, ATP- and metallo-peptidase activity, photoinhibition and PSII repair and catabolic processes; ATP-dependent peptidase activity and PUA-like domain; and transmembrane components.

**Figure 2.**
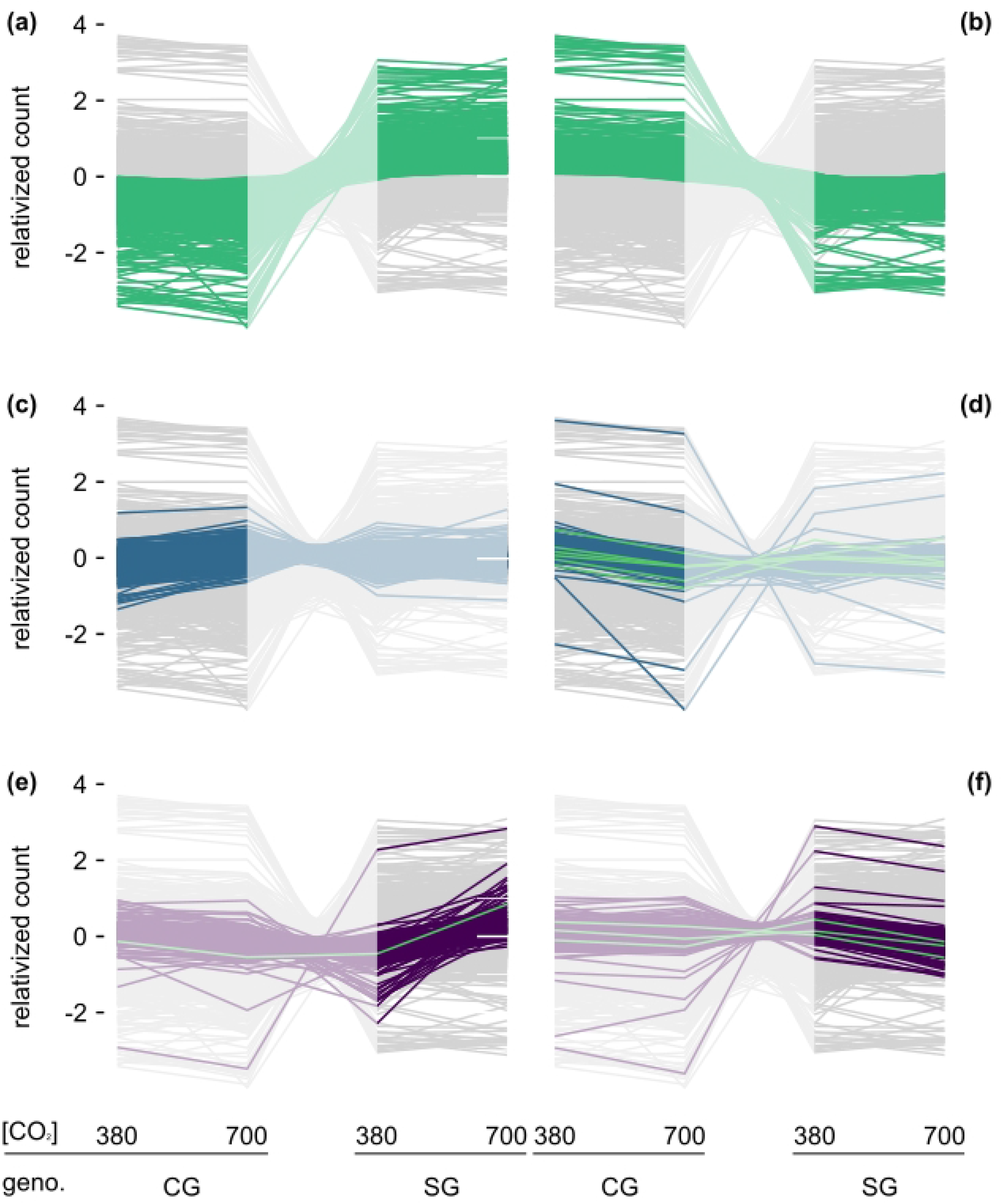
Transcript count differed between the Control and Selected Genotypes (CG and SG) and within genotypes across 380 and 700 ppm [CO_2_]. **(a-f)** Each grey line (background) represents the mean relativized count (centered log ratio) of the 16,472 unique transcript identifiers in this dataset. Colored lines in **bold** represent those transcript identifiers with effect sizes greater than ± 1 in each comparison, while the lighter sections in each plot allow visualization of how those same transcripts respond across genotypes or within the other genotype. **(a-b)** Transcript identifiers showing an increase **(a)** and decrease **(b)** from CG to SG. **(c-d, bold lines)** Transcript identifiers showing an increase **(c)** and decrease **(d)** in CG from 380 to 700 ppm [CO_2_]. For reference, the same transcripts are visible in SG (light lines). **(e-f, bold lines)** Transcript identifiers showing an increase **(e)** and decrease **(f)** in SG from 380 to 700 ppm [CO_2_]. For reference, the same transcripts are visible in CG (light lines). Green lines in **d-f** show flowering genes found to have effect sizes greater than ± 1.

**Table 2:** Summary table of top five functional annotation clusters in each comparison.

| Comparison | Functional Annotation Clusters | Enrich. Score | Unique gene IDs |
| --- | --- | --- | --- |
| **Genotype**<br>*Increase from CG to SG* | Serine/threonine and protein kinase activity | 21.43 | 1573 |
|  | Transmembrane or membrane components | 11.93 |  |
|  | Calcium-binding region, serine/threonine kinase, peptidyl-serine phosphorylation signal transduction | 7.5 |  |
|  | ADP and DNA binding, leucine-rich repeats | 5.68 |  |
|  | Ankyrin repeat and PGG domains | 5.55 |  |
| **Genotype**<br>*Decrease from CG to SG* | Chloroplast | 172.46 | 1987 |
|  | Ribosome | 37.42 |  |
|  | Chloroplast thylakoid lumen | 19.91 |  |
|  | Ribosomal RNA-binding | 9.38 |  |
|  | Lipid biosynthesis and metabolism | 6.11 |  |
| **CG**<br>*Increase from 380 to 700 ppm* | Transit peptide, chloroplast thylakoid membrane | 20.13 | 701 |
|  | PSI, PSII, chlorophyll a/b, chloroplast, magnesium binding | 12.17 |  |
|  | Membrane, transmembrane | 5.89 |  |
|  | Glycolysis/gluconeogenesis, biosynthesis of amino acids/antibiotics | 4.38 |  |
|  | Cytochrome b5 heme-binding | 3.87 |  |
| **CG**<br>*Decrease from 380 to 700 ppm* | ATP and nucleotide binding | 4.93 | 292 |
|  | Microtubule motor protein activity | 4.39 |  |
|  | Nucleus, sequence-specific DNA binding, and transcription regulation | 4.35 |  |
|  | Cell division and mitosis | 3.55 |  |
|  | Zinc and metal binding | 2.76 |  |
| **SG**<br>*Increase from 380 to 700 ppm* | Golgi related and glycosyl and hexosyl transferase activity | 3.75 | 226 |
|  | Small GTP binding and GTPase-mediated signal transduction | 2.83 |  |
|  | Cytoskeleton and microtubule | 2.71 |  |
|  | Cell wall organization | 2.24 |  |
|  | IQ motif and calmodulin binding | 1.79 |  |
| **SG**<br>*Decrease from 380 to 700 ppm* | Chloroplast and transit peptide | 82.03 | 1017 |
|  | Carbon metabolism and fixation, biosynthesis and metabolic pathways | 5.42 |  |
|  | Thylakoid, ATP- and metallo-peptidase activity, photoinhibition and PSII repair and catabolic processes | 4.92 |  |
|  | ATP-dependent peptidase activity and PUA-like domain | 4.55 |  |
|  | Transmembrane components | 4.21 |  |

Many of the significant clusters for each comparison had overlapping annotation categories (**Table 2, S2a- h**), so to better characterize overarching functional descriptions we used both manual categorization and word-cloud generation (worditout.com) to assess patterns across all significant annotation clusters. These categorization processes revealed that there was an increase from CG to SG, relative to the internal control, in serine/threonine protein phosphorylation and signaling, glycoprotein and glycosylation signaling, and Golgi and vesicle transport.

Thus, the two genotypes have different signaling cascades at the developmental stage just prior to flowering.

Conversely, there was a decrease from CG to SG in photosynthetic and energy transfer (redox) processes. This appeared most strongly driven by down-regulation of photosynthesis related genes in SG from 380 to 700 ppm as CG and SG displayed nearly opposite patterns of gene regulation from 380 to 700 ppm. For instance, CG increased genes involved in processes related to photosynthesis, energy transfer, and metabolite breakdown and biosynthesis. Thus, CG may capitalize on available carbon in a high [CO_2_] environment by increasing carbon acquisition and processing. Conversely, SG decreased genes involved in processes related to PSII repair and photoinhibition, energy transfer (i.e. FAD/NADPH), and carbon fixation and metabolite biosynthesis. Perhaps, under current [CO_2_], SG experiences some photoinhibition which is alleviated by increased CO_2_ availability as has been shown for other species when nutrients are not limiting [54,55]. Further, while CG decreased genes involved in processes related to nucleotide binding, cell division, and cell reorganization and internal transport (i.e. motor proteins and nuclear pores), SG increased genes involved in processes related to cellular internal transport (i.e. motor proteins and Golgi vesicles), and sugar signaling and processing (i.e. glycosyl transferase, polysaccharide binding). Thus, when SG is not preparing to flower at elevated [CO_2_], it is maintaining relatively higher intercellular signaling and motility. It is unknown from our current design whether such behavior would be observed in SG grown at 380 ppm well before flowering is initiated.

## Control and Selected Genotypes display differing metabolic responses to elevated [CO_2_]

As CG showed no clear [CO_2_] response in primary carbohydrate levels yet displayed strong changes in carbohydrate-related genes, we wondered whether other metabolites were being altered in CG. We assessed 15 additional metabolites common across the NMR and GCMS data sets (**Table S3**), assessing the effect of [CO_2_] independently within each genotype as the effect of genotype was very strong, and incorporating dataset as a covariate. We noted differing carbohydrate profiles for each genotype, with CG showing significant increases in glucose-6-phosphate, succinic acid, and trehalose, and decreases in aspartic acid, glycine, and threonine. In addition to increases in glucose and fructose, SG showed increases in succinic acid and trehalose, and decreases in glutamine and serine (**Table S3**). Thus, while CG does not show a strong response to [CO_2_] in primary carbohydrates glucose and fructose, CG does display an altered carbohydrate profile in response to [CO_2_] change, although this does not correlate with altered flowering in this genotype.

### Genes involved in sugar modifications and one-carbon metabolism altered in Selected Genotype

To better assess potential pathways contributing to delayed flowering in SG in response to elevated [CO_2_], we explored genes within the carbohydrate-related functional annotation clusters in more detail. We did this through a manual referencing of genes in significant clusters to The Arabidopsis Information Resource (TAIR, [56]), and by searching the clusters for genes involved in the trehalose-6-phosphate, O-GlcNAcylation, and one-carbon metabolism pathways (**Table 3**). SG showed an increase in several genes encoding O-glycosyl hydrolases including AT5G55180, AT3G55430, and AT5G08000 (also called *E13L3*) (**Table 3**). Per TAIR [56], these genes enable cleavage at internal 1,3-beta-D-glucose linkages (endo-1,3-beta glucosidases) to cause the formation of oligosaccharides. It is possible that the cleaved oligosaccharides participate in a signal cascade serving as secondary modifications to protein- or lipid-based molecules. For instance, several sugar transferases were also positively enriched in SG plants grown at 700 ppm [CO_2_]. These included AT3G21190 (ATMSR1), an O-fucosyltransferase; AT2G28080, a UDP-glycosyltransferase; and AT3G58790 (GAUT15), a galacturonosyltransferase, among others (**Table 3**). We also noted that genes associated with one-carbon metabolism, which involves the addition or removal of single carbon units and which is involved in a range of metabolic, epigenetic, or transcriptional regulatory processes [37,57], displayed differences across groups. We focused on genes involved in the process of methylation, which is involved in transcriptional and epigenetic regulatory processes [36,58]. Several methyltransferases were included in this data set and two of these—AT4G37930 and AT5G13050—declined in SG in response to elevated [CO_2_] (**Table 3**). Several genes act upstream in this pathway to generate S-adenosyl-Met ([59] in [57]) which serves as a methyl group donor for methyltransferase reactions. These are cystathionine gamma-synthase (*CGS*), cystathionine beta-lyase (*CBS*), and methionine synthase; all of which are located in the chloroplast [57] explaining their presence in clusters associated with that term (**Table S2h**). *CGS* (AT3G01120) was lower in SG relative to CG, as was a threonine synthase (*MTO1*, AT3G01120). Additionally, *MTO2* showed a within genotype effect to [CO_2_] as it decreased in SG in elevated [CO_2_] relative to current [CO_2_] conditions (**Table 3, S1**). Threonine synthase competes with *CGS* for the substrate *O-*phosphohomo-Ser (OPH) [57]. As *MTO2* decreases in response to elevated [CO_2_] in SG, it is possible that OPH is being used to generate S-adenosyl-met for methyltransferase reactions, despite both *CGS* and *MTO1* being relatively lower in SG than in CG (**Table S1**). Thus, increased production of both oligosaccharides and single-carbon molecules may occur in SG, correlating with a delay in flowering in elevated [CO_2_]. However, as two methyltransferases decreased in SG while several sugar transferases increase, signaling pathways involving oligosaccharides may contribute to the flowering delay.

**Table 3:**
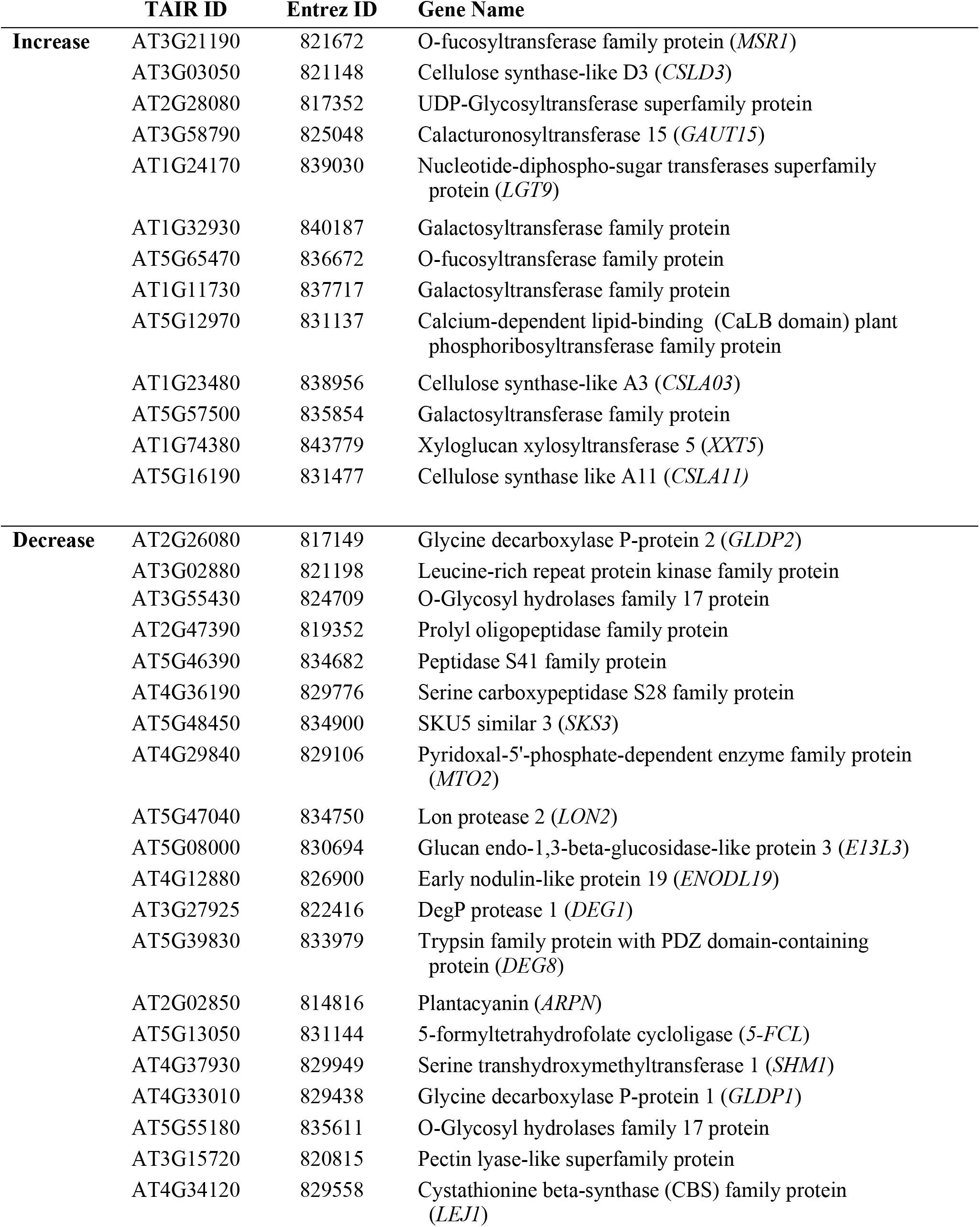
Details of genes found within functional annotations associated with carbohydrate-related processes.

We also determined whether genes from two other carbohydrate-related pathways and associated with flowering differed in this dataset. For genes influencing trehalose-6-phosephate (T6P) [25], eight of the eleven *T6P SYNTHASE* (*TPS*) genes and six of the ten *T6P PHOSPHATASE* genes (reviewed in [60]) were present in this dataset. However, while two T6P-related genes differed between genotypes, only one showed a within-strain [CO_2_] response (**Table S1**). Specifically, AT1G23870 (*TPS9*) decreased from ambient to elevated [CO_2_] in CG; while two members of the ten-member *T6P PHOSPHATASE* family—AT5G65140 (*TPPJ*) and AT2G22190 (*TPPE*)— decreased and increased in SG relative to CG, respectively. Thus, differences in *T6P*-related genes were not observed in SG; although it is important to note that *TPS2, TPS3, TPS4*, *TPPC*, *TPPE*, *TPPG*, and *TPPI* were not included in the dataset. *TREHELASE* (*TRE1*) [61], also included in the dataset, did not show an effect. The T6P pathways interacts with the pathway involving *SUCROSE NON-FERMENTING1-RELATED KINASES* (*SnRK1*, also *KIN10*, AT3G01090) [62], which also influences flowering [63]. While both *KIN10* and related *KIN11* (AT3G29160) [64] were higher in SG than in CG, neither was affected by [CO_2_] change within a genotype (**Table S1**). Secondly, we assessed the genes involved in serine/threonine-linked glycosylation, O-GlcNAcylation [31].

*SECRET AGENT* (*SEC*, AT3G04240) the primary O-GlcNAc transferase in *Arabidopsis*, did not show a large effect size in any of the comparisons made in this work. *SPINDLY* (*SPY*, AT3G11540), an O-fucosyltransferase originally predicted as an O-GlcNAc transferase and which acts within the same regulatory pathways [30,65–67], also did not differ. Thus, these pathways appear not to correspond to the flowering delay in SG.

### Carbohydrate-responsive flowering regulator genes differ in response to [CO_2_] change

The observed differences between genotypes and between the [CO_2_] treatments, primarily in photosynthetic and carbohydrate related processes, are consistent both with the differences in primary carbohydrates we observed, and with the range of photosynthetic and carbohydrate responses observed elsewhere [17]. However, as we were interested in mechanisms controlling flowering, and whether there was a relationship between carbohydrate-mediated pathways and flowering-control mechanisms, we also assessed whether there were differences in the flowering genes independently. We assessed a list of 156 genes known to be associated with flowering [68] (**Table S4**), of which 125 matched transcripts in this dataset (**Table S4**). These included components of the circadian clock, photoperiod, ambient temperature, vernalization, endogenous, and meristem identity flowering control pathways. In CG, there were six genes with effect sizes greater than ± 1, all showing a relative decrease from current to elevated [CO_2_]. These were *ARABIDOPSIS TRITHORAX 2* (*ATX2*); *FRIGIDA* (*FRI*); *AGAMOUS-LIKE 24* (*AGL24*); *GATA, NITRATE-INDUCIBLE, CARBON METABOLISM INVOLVED* (*GNC*); *FRUITFULL* (*FUL*); and *TIMING OF CAB EXPRESSION 1* (*TOC1*). Both *ATX2* and *FRI* regulate the vernalization-responsive, flowering repressor *FLOWERING LOCUS C* (*FLC*), which was previously shown to be strongly elevated in SG under elevated [CO_2_] [15,50], and slightly elevated in CG early in development before declining prior to flowering in a subsequent experiment [50]. *ATX2* is a set-domain-containing protein required for H3K4 methylation and activation of *FLC* [69], while *FRI* complexes with transcriptional and chromatin-modifying factors to induce *FLC* [70]. The MADS-box encoding *AGL24* and *FUL* are floral meristem identity genes downstream of *FLC* and another MADS-box encoding floral regulator gene *SUPPRESSOR OF OVEREXPRESSION OF CONSTANS 1* (*SOC1*) [68], which was previously shown to be suppressed in SG in response to elevated [CO_2_] [15,50] and which was upregulated in SG in elevated [CO_2_] in response to prolonged cold temperatures (vernalization) [50]. *SOC1* is also downstream of *FLC*, which is well known to regulate response to winter temperatures and is repressed by vernalization [71].

Vernalization restored earlier flowering in SG in elevated [CO_2_] [50]. *AGL24* interacts with *SOC1* to regulate another meristem identity gene, *LEAFY* (*LFY*) [72], which was previously shown to be upregulated by elevated [CO_2_] in both genotypes either independently or interactively with prolonged cold treatment (vernalization) [15,50]. *LFY* and *FUL* are regulated by micro RNA 156-regulated SPL transcription factors [73,74], which in turn, are modified by T6P [25], suggesting one mechanism through which [CO_2_]-induced carbohydrate changes may alter flowering. Additionally, *GNC*, known to be involved in stomatal and chloroplast development, glucose sensitivity, and leaf starch level [75–78], regulates *SOC1* and flowering [79,80]. Finally, *TOC1*, is a component of the *Arabidopsis* circadian clock [81], which acts through the photoperiod sensing pathway to regulate flowering time [82,83]. The circadian clock regulates plant metabolic state by regulation of photosynthesis and starch breakdown [84,85] but is carbohydrate responsive as well [86,87]. Thus, although CG does not visibly alter its flowering time in response to [CO_2_], several flowering-regulator genes are altered, many of which have known links to carbohydrate response pathways and to previous responses to [CO_2_].

In SG, there were four genes with effect sizes greater than ± 1, with three showing a relative decrease from current to elevated [CO_2_] and one showing an increase. Those showing a decrease were *SUPPRESSOR OF PHYA- 105 3* (*SPA3*), *BOTRYTIS SUSCEPTIBLE 1 INTERACTOR* (*BIO*), and *CONSTANS* (*CO*). The one showing an increase was *TEMPRANILLO 2* (*TEM2*). *CO* is a key, circadian-regulated component of the photoperiodic-sensing pathway and upstream inducer of the floral integrator gene, *FLOWERING LOCUS T* (*FT*) [88–90]. The SPA1, SPA3, and SPA4 proteins redundantly degrade CO protein in the dark, ensuring delayed flowering in short-photoperiod conditions [91–94]. *BIO* represses flowering by repressing *FT* through both *CO*-dependent and - independent mechanisms [95]. *TEM2*, along with *TEM1*, acts to repress *FT* antagonistically to CO, with TEM1 recruiting Polycomb factors to the *FT* locus [96,97]. Both *BIO* and the *TEM* genes are associated with the gibberellin pathway [98,99], and several members of the gibberellin-response pathway appear also to be involved in plant carbohydrate regulation and flowering time [100]. Thus, across both genotypes, genes within the photoperiod, vernalization, and meristem identity response pathways are altered in response to [CO_2_].

## Discussion

Here, we aimed to assess potential metabolic pathways through which [CO_2_] change may alter flowering genes and flowering times and to determine mechanisms for genotypic variation in flowering response. We found that prior to flowering, the *Arabidopsis* Selected Genotype (SG) that delays flowering when exposed to elevated [CO_2_] had higher sucrose levels relative to the Control Genotype (CG) that does not show a [CO_2_]-induced flowering phenotype. SG also responds more strongly than CG to a [CO_2_] increase by increasing glucose and fructose levels. This pattern is consistent with the delay in flowering in *Arabidopsis* observed over increasing sucrose concentrations in plant growth media [24]. Thus, higher sugar content in the form of glucose and fructose may contribute to the flowering delay in SG in response to elevated [CO_2_]. *Arabidopsis* accessions display substantial variation in carbon accumulation and photosynthetic capacity in general [101] and the degree to which these traits respond to [CO_2_] and temperature change [45,102]. As SG appears to respond more strongly to [CO_2_] change by altering these foliar carbohydrates, capacity to accumulate higher glucose and fructose could be an indicator of flowering time change in response to [CO_2_] rise. Finally, it is also possible that higher sucrose accumulation as observed in SG indicates likelihood to delay flowering. However, while both CG and SG were collected at analogous developmental stages (i.e. before the reproductive transition), SG was collected at a higher leaf number. Thus, subsequent studies exploring change in carbohydrate accumulation over time will be useful, as will experiments assessing whether these patterns hold across a broad range of accessions.

A large body of work now demonstrates the connection between transcriptional and epigenetic regulatory processes and carbohydrate levels in both plants and animals (as reviewed in [36,103]). Small soluble sugars such as glucose and fructose serve direct signaling functions [103], but carbohydrates also serve as protein post-translational modifications and as substrates for protein and histone methylation [36,103]. In animals, these processes link the nutrition or disease state of an organism to the genome to elicit a response [26,104]. In plants, these processes likely link not only the endogenous environment but the external environment to the genome as carbohydrate levels in plant tissue are altered by cold temperatures [105,106], drought [107,108], salinity [109,110], daylength and light levels [20,111], and herbivory [112] to name only a few environmental variables. Thus, it stands to reason that changes in carbohydrate composition and level would be the mediating factor through which change in atmospheric [CO_2_], the primary photosynthetic substrate, alters developmental responses such as the timing of the vegetative to reproductive transition. Here, during the developmental stage prior to flowering, both CG and SG displayed alterations in genes related to metabolic processes in response to elevated [CO_2_] relative to current [CO_2_]. However, the two genotypes differed in the processes altered. CG increased genes related to breakdown of fructose 1,6- bisphosphate (fructose-bisphosphate aldolase) and the synthesis of amino acids and secondary metabolites.

Conversely, in response to a rise in [CO_2_], SG showed a decrease in genes related to starch and sugar biosynthesis and metabolism, but increased genes involved in production of oligosaccharides and in sugar modifications (glycosyl and hexosyl transferase, polysaccharide binding). Additionally, one of the threonine synthases (*MTO2*) present in the data set decreased in response to elevated [CO_2_] in SG. These enzymes competitively inhibit reactions involved in the production of S-adenosyl-Met, which donates methyl groups for methyltransferase reactions [57].

Thus, increases in oligosaccharide pools and potentially the pool of S-adenosyl-Met correlate with delayed flowering in SG in response to elevated [CO_2_]. The oligosaccharide pool is likely contributing to a signaling cascade as glycosyl and hexosyl transferases also increase. However, as small sugars can act independently or modify both proteins and lipids [103], whether the oligosaccharide increase is influencing specific pathways or acting more generally is an open question.

Finally, as we were interested in the flowering regulatory mechanisms correlating with differences in [CO_2_] response in these genotypes, we explored flowering-related genes specifically and noted that several genes representing the vernalization, photoperiod, and meristem identity pathways were altered in either CG or SG in response to [CO_2_] change. Although CG shows no flowering time phenotype in response to elevated [CO_2_], flowering genes *FLC*, *SOC1*, and *LFY* were shown to be altered in previous studies using these lines, and genes associated with all three were altered in CG here. We noted differences in *ATX2* which appears to act semi-redundantly with *ATX1* to regulate *FLC* expression [113], but may play more of a role later in development as *pATX2:GUS* was expressed in older leaves while *pATX1:GUS* was expressed throughout development [114].

Although, to our knowledge, *ATX2* has not yet been associated with carbohydrate changes, *ATX1* is regulated by O-GlcNAcylation [31] and *ATX5* is glucose responsive [115]. Thus, *ATX2* may respond to [CO_2_]-induced foliar carbohydrate changes to influenced *FLC* at least in CG. We also noted alterations in *AGL24* and *GNC* which interact with *SOC1*, a gene upstream of *LFY* [72,80]. Here, we noted alterations in *FUL* as well. Per the Flowering Interactive Database, *LFY* and *FUL* are both regulated by *SOC1* [68], but are also regulated by *SPL* transcription factors which are influenced by T6P [25,73,74]. Although T6P-related genes were not altered in SG in response to [CO_2_] change, *TPS9* was altered in CG. *TPS9* seems not to act enzymatically to affect *T6P* levels, but may serve a regulatory function in response to carbohydrates and is repressed by sucrose and glucose [116] consistent with its decrease from current to elevated [CO_2_], here. While we did not observe altered sucrose, glucose, or fructose in response to [CO_2_] change in CG, other carbohydrates did increase with elevated [CO_2_] in that genotype including glucose-6-phosphate and trehalose, perhaps leading to the changes observed here. Finally, the circadian clock gene *TOC1* regulates the photoperiod-sensing pathway upstream of *SOC1*, *LFY*, and *FUL* [68]. The circadian clock is also regulated by carbohydrates [86,87], demonstrating that [CO_2_] rise may influence the expression profiles of several flowering-related pathways.

Although the vernalization response pathway in addition to other floral integrator genes has been shown to be important for the [CO_2_]-induced delay in SG [15,50], we did not note vernalization response genes to be altered here in SG. However, we noted that genes involved in the photoperiod and gibberellin-response pathways were altered. These genes all act upstream of the key floral integrator genes, *FT*, *LFY*, and *SOC1*, and also have connections to carbohydrates. For instance, the SPAs contain a serine/threonine protein kinase domain that, in SPA1, has recently been shown to be necessary for photomorphogenic response [117]. Protein phosphorylation and the protein sugar modification, O-GlcNAcylation, both target serine and threonine amino acids and are known to both compete for and influence the other [118]. Additionally, the SPAs redundantly complex with *CONSTITUTIVE PHOTOMORPHOGENIC 1* (*COP1*) to degrade CO protein in the dark [91–94]. PHYTOCHROME A disrupts the COP1/SPA complex to allow light-promoted flowering; however, COP1/SPA may feedback to degrade PHYA in a manner that is dependent on sugar [119,120]. Additionally, *CO*, *COP1*, and the *SPA* family are all regulated by the circadian clock, a process that is also influenced by carbohydrates [86,87]. Additionally, both *BIO* and *TEM* genes are involved in the gibberellic (GA) sensing pathway, which is associated with sugar sensing at several points. For instance, GA and sucrose were implicated early in their interactive activation of *LFY* [121], and the *TEM* genes directly regulate genes involved in GA biosynthesis as well as influence the expression of *FT* which acts upstream of *LFY* and other flowering-transition genes [68,99]. Further, BIO and DELLA proteins interact to regulate GA-responsive genes [98], while *GIGANTEA* (*GI*), a key circadian clock gene, stabilizes the DELLA proteins to gate GA-response to the night [122]. The DELLAs are post-translationally modified by O-GlcNAcylation, while *GIGANTEA* is involved in sugar-sensing and in sugar regulation of the circadian clock [123]. Whether *BIO* and *TEM2* transcription is influenced either directly or indirectly by sugars is, to our knowledge, not known. However, in rice, sugar starvation directly influenced a regulator of genes involved in GA biosynthesis [124]. Thus, across CG and SG, our study highlights several additional candidate flowering-control pathways responsible for flowering time variation in response to [CO_2_].

It should be noted that while we had observed the key vernalization-responsive, floral-repressor gene, *FLC*, to prolong its elevated expression in SG and that its expression contributed to flowering delays [15,50], *FLC* did not show differences in either CG or SG at the time point harvested, here. This may be due to the detection method used as we noted significant variation among samples. Additionally, the flowering genes, overall, showed a much lower degree of response than genes related to carbohydrate pathways and therefore differences detected by exploring these genes individually may not be detected through a transcriptomics approach. Further, while several genes upstream of key floral integrators *FT* and *LFY* were altered in our dataset, neither *FT* nor *LFY* were included in the dataset. Finally, regulation of these genes is complex, with several cycling over a 24-hour period as regulated by the circadian clock [68] or changing throughout development as is the case of vernalization-responsive genes [34].

Thus, much more work needs to be done to understand the relative influence of the different flowering pathways shown to be altered here and the degree to which they interact. Further experiments assessing their degree of change over developmental and diurnal time will be necessary.

In sum, this study coupled with our two previous studies paints a picture in which atmospheric [CO_2_] change influences carbohydrate response pathways, which in turn influence flowering regulators to alter flowering in a genotype-specific manner. This study highlights additional candidate pathways responsible for flowering variation in response to atmospheric [CO_2_] change.

## Materials and Methods

### Plant Material and Growth Conditions

We used our novel *Arabidopsis* system involving two genotypes [14,15,125], whereby genotype SG delays flowering at elevated [CO_2_] and flowers at a larger size, and genotype CG exhibits similar flowering times and size at flowering between 380 and 700 ppm [CO_2_]. These genotypes were originally generated from the same parental cross. SG was generated through selection over consecutive generations of growth at elevated [CO_2_], by choosing individuals with high seed set [14]. CG was generated through selection of random individuals at each generation. We assessed carbohydrate, metabolite, and transcriptomic responses in rosettes leading up to flowering in SG and CG plants grown at 380 and 700 ppm [CO_2_] (**Fig. 1**). For SG, these measurements were taken prior to the initiation of reproduction at 380 ppm [CO_2_] as well as the analogous stage (leaf number) during growth at 700 ppm when plants should flower (as in 380 ppm), but do not (see **Fig. 1** for harvesting regime). Sample size (*n*) was five to nine plants per genotype for both metabolomic and transcriptomic analyses. Flowering was defined as the visible transition from vegetative to floral growth of the meristem (i.e. the flowering stem was visible above the rosette).

Signal plants were planted out one week prior to the plants used in the experiment, such that when the signal plants visibly transitioned to reproduction, the plants used for the experiment were harvested. Rosette leaf numbers were counted at harvest and cotyledons were excluded from these counts.

We utilized Conviron BDR16 (Winnepeg, Canada) growth chambers with custom control of [CO_2_], in which [CO_2_] was automatically injected when needed and chamber air was pulled through JorVet soda lime (Loveland, CO, U.S.A.) to scrub excess [CO_2_]. Chambers constantly monitored internal conditions and [CO_2_] was maintained at ±20 ppm of either 380 or 700 ppm at least 95% of the time. Temperatures were set at 25/18 °C day/night and humidity was set at 60/90% day night. Seeds were cold stratified at 4 °C for four days prior to beginning the experiment to promote uniform germination. Plants were grown under 14-hour photoperiods with light levels ∼800 µmol m-2 s-1 in 750 mL pots filled with a 1:1:1 (v/v) mixture of pea gravel, vermiculate, and Terface (Profile Products, Buffalo Grove, IL, U.S.A.). All plants were well watered and dosed with half-strength Hoagland’s Solution (**Table S5**) daily.

### Carbohydrate and Metabolite Profiling

In collaboration with the Ecological and Molecular Sciences Laboratory (EMSL) at the Department of Energy Pacific Northwest National Laboratory EMSL, we measured total leaf sucrose, glucose, fructose and related metabolites during time points leading up to flowering in SG and CG plants grown at 380 and 700 ppm [CO_2_]. For NMR, *Arabidopsis thaliana* frozen leaf tissues were weighed and ground by using two 3 mm stainless steel beads for 3 minutes at 30 Hz with frozen adapters on a TissueLyser II (Qiagen). The resulting frozen powder was dissolved in 650 µL chloroform-methanol (3:7, v/v) and placed in the -20 °C freezer with occasional shaking for 2 hours. Next, 600 µL of ice-cold nanopure water was added and placed in the 4 °C fridge with repeated shaking for 15 minutes. Finally, the sample was centrifuged at 12,000 rpm for 5 mins, the aqueous phase was collected and dried in the speed-vacuum concentrator. The NMR sample of *Arabidopsis* was dissolved in 600 µL of H_2_O-D_2_O (9:1,v/v) with 0.5 mM DSS.

All NMR spectra were collected using a Varian Direct Drive 600 MHz NMR spectrometer equipped with a 5 mm triple-resonance salt-tolerant cold probe. The 1D 1H NMR spectra of all samples were processed, assigned, and analyzed by using Chenomx NMR Suite 8.1 with quantification based on spectral intensities relative to the internal standard. Candidate metabolites present in each complex mixture were determined by matching the chemical shift, J-coupling, and intensity information of experimental NMR signals against the NMR signals of standard metabolites in the Chenomx library. The 1D 1H spectra were collected following standard Chenomx data collection guidelines [126], employing a 1D NOESY presaturation experiment with 65536 complex points and at least 512 scans at 298 K. Additionally, 2D ^13^C-^1^H HSQC spectra were collected with N1=1024 and N2=1024 complex points. The spectral widths along the indirect and direct dimension were 160 and 12 ppm, respectively.

The number of scans per t_1_ increment was 16. 2D ^1^H-^1^H TOCSY spectra of *Arabidopsis thailana* metabolite extract were collected with N1=512 and N2=1024 complex points. The spectral widths along the indirect and direct dimension were 12 ppm and TOCSY mixing time was 90 ms. 2D spectra (including ^1^H-^13^C heteronuclear single-quantum correlation spectroscopy (HSQC),^1^H-^1^H total correlation spectroscopy (TOCSY)) were acquired on most of the leaf extract samples, aiding as needed in the 1D ^1^H assignments.

Gas chromatography-mass spectrometry (GC-MS) based untargeted analysis of extracted metabolites was following Xu and colleagues [127]. The polar metabolites were completely dried under speed vacuum concentrator, then, chemically derivatized and analyzed by GC-MS. Metabolites were derivatized as previously described [128] by adding 20 µl of methoxyamine solution (30 mg/ml in pyridine) and incubated at 37 °C for 90 mins. to protect the carbonyl groups and reduce carbohydrate isoforms. Then 80 µl of N-methyl-N-(trimethylsilyl)-trifluoroacetamide with 1% trimethylchlorosilane were added to each sample to trimethylsilyate the hydroxyl and amine groups for 30 mins. The samples were cooled to room temperature prior to GC-MS analysis. Data collected by GC-MS were processed using the MetaboliteDetector software, version 2.5 beta [129]. Retention indices of detected metabolites were calculated based on analysis of the fatty acid methyl esters mixture (C8 - C28), followed by chromatographic alignment across all analyses after deconvolution. Metabolites were initially identified by matching experimental spectra to a PNNL augmented version of the Agilent Fiehn Metabolomics Library containing spectra and validated retention indices for over 900 metabolites [130], and additionally cross-checked by matching with NIST17 GC-MS Spectral Library. All metabolite identifications were manually validated to minimize deconvolution and identification errors during the automated data processing.

The NMR and GCMS datasets were conducted on separate experimental replicates, which displayed variation in their overall responses that we attributed to effect of replicate. To account for this, we analyzed only those metabolites in common between the two datasets (**Table S3**), first transforming the data using centered-log ratio [51,52], then using Analysis of Variance (ANOVA, function *aov* in R v. 4.1.1) considering the effects of genotype, [CO_2_], their interaction, and experimental replicate as a covariate. Each metabolite was analyzed separately, then the *p-*values adjusted for multiple comparisons using the *p.adjust* function in base R (v. 4.1.1, method = fdr) [131]. *Post hoc* analysis through Tukey’s Honest Significant Difference (function *TukeyHSD*, v. 4.1.1), was used to assess differences among treatment groups.

### Transcriptomic Profiling

#### a. Assembly of the SG and CG genomes

RNAseq and related bioinformatics were conducted in partnership with EMSL utilizing reference genomes for SG and CG sequenced through the University of Kansas Genomics CORE facility. Genomic DNA isolation, library preparation, and sequencing were as previously described [50]. Specifically, genomic DNA was isolated using the DNeasy Plant kit (Qiagen, Denmark) from two pooled, fully inbred plants from both the CG and SG lines. The libraries were prepared using the TruSeq DNA PCR-Free kit and sequenced on the HiSeq RR-PE100 system (Illumina, USA). This resulted in approximately 188 million reads in total or about 94 million reads per pooled sample, and about 200x coverage. These 100-bp reads were assembled into genes-only CG and SG genomes using ABySS assembly software (version 1.9.0, doi: <u>10.1101/gr.214346.116</u>) using K=96. To predict the location of genes on the assembled sequences, we used the gene calling web server Augustus (http://bioinf.unigreifswald.de/augustus/submission.php). For each sequence predicted to be a gene, gene annotation was acquired using the best hit from a BLASTP search, using the plant component of Uniprot combined with the Araport11 gene set (https://www.araport.org/data/araport11). Genes received a genome specific identifier, as well as were matched with their most closely corresponding locus-linked *Arabidopsis* gene ID.

#### b. RNA sequencing

RNA was isolated using Qiagen RNeasy™ mini kit (cat#74104), followed by genomic DNA removal and cleaning using Qiagen RNase-Free DNase Set kit (cat#79254) and Qiagen Mini RNeasy™ kit (cat#74104) (Qiagen, Denmark). Integrity of the RNA samples was assessed using the Alegient 2100 Bioanalyzer. RNA samples having an RNA Integrity Number between nine and ten were used in this work. rRNA was removed using Ribo-Zero rRNA removal kit (cat#MRRZPL1224, Illumina, San Diego, CA, USA). The SOLiD™ Total RNA-Seq Kit (cat#4445374) was used to construct template cDNA for RNASeq following the whole transcriptome protocol recommended by Applied Biosystems. Briefly, mRNA was fragmented using chemical hydrolysis followed by ligation with strand-specific adapters and reverse transcript to generate cDNA. The cDNA fragments, 150 to 250 bp in size, were isolated and amplified through 15 amplification cycles to produce the required number of templates for the SOLiD™ EZ Bead™ system, which was used to generate the strand-specific template bead library for ligation base sequencing by the 5500xl SOLiD™ instrument (LifeTechnologies, ThermoFisher, Carlsbad, CA, USA). The 50-base short read sequences produced by the SOLiD sequencer were mapped in color space using the Whole Transcriptome analysis pipeline in the Life Technologies LifeScope software version 2.5 against the genes-only genomes assembled, as described above, for the CG and SG Arabidopsis strains as well as the Araport11 reference genome.

#### c. Transcriptome analysis

Alignments to genotype-specific and Araport11 reference genomes were compared, and the Araport11 alignments were selected for continued use, as read count was higher. Specifically, count datasets for individual samples were compiled into a single dataset using R (v. 3.6.3), and rows containing duplicate gene identifiers were averaged for each individual as counts were similar or the same across rows containing the same gene identifiers (using the *aggregate* function in R). This resulted in 31,556 unique transcript identifiers. Individuals and counts were assessed for quality using the *goodSamplesGenes* function in the *WGCNA* package in R (v. 3.6.3, minFraction = ¾), then transcript identifiers containing mostly zeros were removed using the *count_filter* function in the *ERSSA* package in R (v. 3.6.3, cutoff = 1). This resulted in 16,472 unique transcript identifiers.

These remaining counts were analyzed using a workflow previously suggested for compositional data (i.e. data for which the upper bounds are limited by detection method and thus not representative of the true high values) [51,52]. Through this method, counts within a sample are relativized against the centered log ratio (clr) of all transcripts within a sample. To do so, any remaining zeros were replaced with very low values using the *cmultRep1* function in the *zCompositions* package in R (v. 4.1.1), then the clr was calculated for each transcript identifier and sample using the *clr* function in the *rgr* package in R (v. 4.1.1). These relativized values were used to calculate the Aitchison distances among samples using the *dist* function in the *robCompositions* package in R (v. 4.1.1, method = euclidean). Samples were clustered based on these distances using the *hclust* function, and two samples, one in CG and one in SG were determined to be outliers as they grouped together, but independently of all other samples in each genotype (**Fig. S1**). These samples were removed, then the distances recalculated. The Aitchison distances were then used to explore relationships among samples and across genotype and [CO_2_] treatments using Principal Component Analysis (PCA, *prcomp* function in R) and multivariate comparison using the *adonis* function in the *vegan* package in R (v. 4.1.1), which allows for interactions among treatments. Once these broad patterns among samples were determined, differential expression analysis of independent transcript identifiers was conducted across genotypes and [CO_2_] treatments (*aldex.clr* function, *ALDEx2* package), and across [CO_2_] treatments within each genotype separately (*aldex* function, *ALDEx2* package). Transcript identifiers with effect sizes greater than ±1 [52] were pulled out for functional annotation analysis using DAVID and the functional annotation clustering function (david.ncifcrf.gov) [132,133], which pulls annotation terms from multiple resources and clusters those terms based on the overlap in genes used to call each term. Clusters with *p* values of 0.05 were considered significant. A list of 156 flowering regulator genes, of which 125 corresponded to the transcript identifiers in this dataset, were analyzed separately for effect sizes calculated between [CO_2_] within a genotype using the following: (*µ*_1_ - *µ*_2_)/mean(*σ*_1_, *σ*_2_), where *µ* and *σ* are the group mean and standard deviation for a transcript identifier across a treatment group.

## Data availability statement

Raw data and codes for analysis will be made available on Open Science Framework at the time of publication. View-only link for review: https://osf.io/zbdq2/?view_only=a117caf2d2e74bc28a3442bba742080c

## Acknowledgements

The authors thank Aleah Henderson-Carter for her help in maintaining and conducting the experiments, Dr. Lena Hileman for her ongoing encouragement, and Drs. John Kelly, Maggie Wagner, and the participants of EEB Genetics for their helpful advice in the analysis. We also thank the University of Kansas Genome Core Facility for their role in this work and their technical support. This work was supported by National Science Foundation (IOS- 1457236) to JKW, and the National Institute of Health Research and Academic Career Development Award (IRADCA) fellowship K12GM063651 to HK-S. A portion of this research was performed on a project award (10.46936/sthm.proj.2013.47814/60005186) from the Environmental Molecular Sciences Laboratory, a Department of Energy Office of Science User Facility sponsored by the Biological and Environmental Research program under Contract No. DE-AC05-76RL01830. Pacific Northwest National Laboratory is a multi-program national laboratory operated by Battelle for the Department of Energy under Contract No. DE-AC05-76RLO1830. The funders has no role in study design, data collection and analysis, decision to publish, or preparation of the manuscript.

## Author Contributions

H.K.S. Conducted much of the data analysis, data interpretation, and manuscript writing.

S.M.W. Designed and conducted the experiment, conducted early data analysis, wrote portions of the manuscript.

K.B. Contributed to experimental design for metabolite profile, conducted NMR analysis, reviewed manuscript.

D.W.H. Contributed to experimental design for metabolite profile, conducted NMR analysis, Chenomx metabolite profiling and develop metabolite workflow, contributed to manuscript.

Y-M.K. Conducted GCMS analysis, contributed to manuscript.

L.M.M. Conducted RNA sequencing and alignments to reference genomes, contributed to manuscript.

H.D.M. Conducted transcript annotations and early transcriptomic comparisons, advised on transcriptomic analyses.

C.D.N. Prepared samples for analyses, reviewed manuscript.

R.T. Conducted genome assembly, contributed to manuscript.

J.K.W. Advised on the design of the experiment and data analysis. Edited and reviewed the manuscript.

## Supplemental figures and tables

Figure S1. Hierarchical cluster analysis of samples based on Aitchison distances calculated using transcript profiles. Outliers shown in red. These were removed for subsequent analyses.

Figure S2. Principal Components Analysis based on Aitchison distances calculated using transcript profiles. Samples grouped by genotype and [CO_2_].

Table S1. List of transcript identifiers having effect sizes greater than ± 1. Transcripts are sorted by increase or decrease from the control, which is either the Control Genotype (CG) for between species comparisons or 380 ppm [CO_2_] for within species comparisons. *(Included as separate tab delimited (.txt) file.)*

Table S2. Full functional annotation tables (outputs from DAVID) for each treatment group stored as .txt files. Table S2a provides is an expanded version of Table 2 in the main text including all Functional Annotation Clusters with Enrichment Scores greater than 1.3. *(All files are included as separate tab delimited (.txt) files.)*

Table S3. ANOVA results of within genotype comparisons of the effect of [CO_2_], incorporating dataset, GCMS or NMR, as a covariate.

Table S4. Flowering genes assessed for effect size in within genotype comparisons of the effect of [CO_2_]. Whether genes were present in transcript dataset is indicated. *(Included as separate tab delimited (.txt) file.)*

Table S5. Hoagland’s Solution.

